# Comparative study of thermal tolerance and other physiological characteristics in *C. briggsae* and *C. elegans*

**DOI:** 10.1101/2023.08.03.551855

**Authors:** Nikita Jhaveri, Harvir Bhullar, Bhagwati P. Gupta

**Affiliations:** Department of Biology, McMaster University, Hamilton, ON L8S 4K1, Canada

**Keywords:** *C. briggsae*, nematode, genetics, comparative study, evolution, reproduction, heat tolerance, thermal resistance, environmental stress

## Abstract

The nematode *Caenorhabditis briggsae* (*C. briggsae*) is routinely used in genetic and evolutionary studies involving its well-known cousin, *C. elegans*. The two species are morphologically almost identical but exhibit significant developmental, genetic, and genomic differences. The AF16 isolate of *C. briggsae* is an established reference strain. We used additional wild isolates from tropical and temperate regions to perform a comparative study of phenotypic characters. The analysis revealed both intra (between *C. briggsae* isolates) and inter (compared to *C. elegans* N2) species variability in dimensions and opacity. Our data also showed that *C. briggsae* isolates prefer higher temperatures for growth, reproduction, and survival than *C. elegans* N2. The preference for higher temperatures further translated into a higher tolerance for heat stress, as evidenced by its survival at temperatures lethal to N2. Interestingly, we found that while *C. briggsae* is more resistant to heat and shows greater sensitivity to other forms of stress, namely oxidative and osmotic, compared to *C. elegans*. The heat resistance of *C. briggsae* was correlated with efficient upregulation of the cytosolic chaperon *hsp-16.2*. Overall, this work has revealed significant differences in stress sensitivities between the two nematodes and forms the basis to investigate changes in underlying mechanisms that affect their stress responses.

## INTRODUCTION

Nematodes of the *Caenorhabditis* genus are used extensively for the in vivo study of biological processes and comparative analysis of gene function. In particular, the nematode *Caenorhabditis elegans* (*C. elegans*) is an established model system for biological research. Since its first use by Sydney Brenner (Brenner, 1974) *C. elegans* has been one of the few chosen organisms to study the genetic basis of animal development, behavior, and physiology (Meneely et al., 2019). Many features of *C. elegans* have contributed to its value as a leading animal model including a short life cycle, inexpensive culture condition, small footprint, transparency, fewer cells, and the ease by which genes can be modified and functionally assayed in the laboratory.

Comparative and evolutionary studies in nematodes have typically involved *C. elegans* and a small set of *Caenorhabditis* species, including *C. briggsae* (for review see Sommer and Bumbarger, 2012). *C. briggsae* was first isolated by Margaret Briggs at Stanford University in Palo Alto, California (Briggs, 1946), who studied the life cycle of animals and also reported living bacteria serving as a food source. (Dougherty and Nigon (1949) recognized the potential of *C. briggsae* for comparative physiology and genetic studies. In fact, Sydney Brenner had initially intended to use *C. briggsae* as his system of choice (Félix, 2008). *C. elegans* and *C. briggsae* diverged from a common ancestor approximately 30 million years ago (Cutter, 2008). Both are androdiecious although their hermaphroditic lifestyle has evolved separately (Kiontke et al., 2004). Similar to *C. elegans*, *C. briggsae* offers many experimental advantages including the ease of cultivation and genetic manipulations in the laboratory (Gupta et al., 2007). The two species also share other features such as gut microbiomes (Dirksen et al., 2016), food preferences (Schulenburg and Félix, 2017), and vector carriers (Kiontke and Sudhaus, 2006). Available genetic and genomic resources for *C. briggsae* include a fully sequenced genome (Stein et al., 2003), chromosome-level assembly (Hillier et al., 2007; Ross et al., 2011) more than 30,000 polymorphisms (Koboldt et al., 2010) that allow mapping of mutants isolated in forward genetic screens (for example, see (Guo et al., 2013; Seetharaman et al., 2010; Sharanya et al., 2015; Jhaveri and Gupta, 2023), operons and 5’ sequence tags (Jhaveri et al., 2022), mutant strains (Gupta et al. in preparation), wild isolates collected from places around the world (Cutter et al., 2006; Félix and Duveau, 2012; Crombie et al., 2019), recombinant inbred lines (Ross et al., 2011; Stevens et al., 2022), proteome analysis (An et al., 2017) similar to that conducted in *C. elegans* (Grün et al., 2014; Tabuse et al., 2005) and an RNAi library covering 1,333 genes (Verster et al., 2014). Morphological and behavioral studies of *C. briggsae* have revealed that while the species is very similar to *C. elegans*, there are significant differences such as excretory duct placement (Wang and Chamberlin, 2002), male tail ray pattern (Fitch, David H.A. Emmons, 1995; Fitch, 1997) and thermal sensitivity (Prasad et al., 2011; Petrella, 2014; Begasse et al., 2015).

In this study, we have further investigated the traits of *C. briggsae* using multiple isolates from tropical (AF16 and QX34) and temperate (HK104 and VX1410) regions. Our results showed that AF16 is significantly smaller and thinner when compared to *C. elegans* N2. However, this was not the case for other *C. briggsae* isolates, suggesting significant variation in dimension across isolates and between species. Likewise, the transparency of animals was variable depending on the isolate. We examined spontaneous males in unmated AF16 hermaphrodite cultures and found a higher frequency compared to N2, which indicated differences in non-disjunction rates between the two strains. Another observation was the altered thickness of the uterine-seam cell (utse) that forms a vulva-uterine connection in hermaphrodites of these species. A significant finding of this work is that *C. briggsae* shows higher resistance to acute heat shock treatment than *C. elegans* and prefers elevated temperatures for growth and reproduction. Interestingly, the animals showed greater sensitivity to two other forms of stress: osmotic and oxidative stress. The heat tolerance phenotype of *C. briggsae* is likely dependent on the cytosolic chaperon *hsp-16*.2 since temperature pulses caused a massive increase in *hsp-16*.2 expression, with levels showing more dynamic changes compared to *C. elegans*. Overall, the findings presented in this study demonstrate significant differences between the two species and provide valuable data to investigate the molecular genetic basis of thermal response in *C. briggsae*.

## MATERIALS AND METHODS

### Strains and culture conditions

*C. briggsae* wild isolates: AF16, HK104, VX34 and QX1410. *C. elegans*: N2. The *C. briggsae* AF16 has been extensively used as a reference strain (Baird and Chamberlin 2006). Worms were grown at 20°C on NG-Agar plates using standard culture conditions (Brenner, 1974) except where specified. The plates were seeded with *Escherichia coli* bacteria (OP50) that served as a food source for worms (Stiernagle, 2006). For Nomarski imaging, worms were mounted on glass slides containing a thin agar pad (2% noble agar). The animals were placed in a drop of M9 buffer that included 5 mM Sodium Azide anesthetic (Yochem, 2006) and examined under the upright Zeiss Zeiss Axioimager D1 and Nikon Eclipse 80i microscopes. Images were captured using Nikon Eclipse 80i and Zeiss Zen 3.0 software.

### Size and movement characteristics

Length and width of anesthetized day-1 adults were measured using Zeiss Zen 3.0 software. For this, animals were mounted on glass slides and examined under the Nomarski microscope. Movement in day-1 adults was analyzed using the tracks in thin overnight grown bacterial lawns on agar plates. Each worm was allowed to move for 30 seconds, and amplitude was measured.

### Frequency of males

To quantify the frequency of males, 4 - 5 L4 hermaphrodites were kept at 15^0^C, 20^0^C and 25^0^C. The worms were allowed to grow till adulthood and lay eggs for one day. Mothers were then removed from the plate. The offspring were screened for males and the male frequency was determined.

### Utse analysis

The thickness of the uterine seam cell (utse) at the top of the vulval apex appearing as hymen was measured in L4 stage larvae. For this, animals were mounted on a glass slide containing agar pad and examining under the Nomarski microscope. Images were captured using Zeiss Zen 3.0 software. The center of the hymen was used for measurements.

### Opacity measurements

Opacity, or optical density, of day-1 adult hermaphrodites was measured by capturing images using a Nomarski microscope and analyzing those using Image J (https://imagej.nih.gov/ij///index.html). Lipids were quantified by the Oil Red O staining method as described previously (Ranawade et al., 2018). Among the techniques used to measure fat levels in *C. elegans*, Oil Red O staining has shown to be the true representation of stored fat content, and positively correlates with the levels of triglycerides (Yen et al., 2010).

### Developmental time analysis

To measure the time required to reach young adults, 4 - 5 gravid hermaphrodites were allowed to lay eggs on a plate for five hours. Mothers were then removed from the plate. The progenies were observed for when they became young adults. Readings were taken every hour after they reached mid L4 stage and were stopped when 80% of the population turned young adults.

### Life span and related physiological characteristics

Life span analysis was carried out as previously described (Amrit et al., 2014). Briefly, 30 - 40 L4 animals were transferred to fresh OP50 plates and kept at 15^0^C, 20^0^C and 28^0^C. The worms were scored every day for survivability. One of the challenges with *C. briggsae* lifespan assay was the tendency of animals to frequently escape the bacterial lawn and commit suicide (most frequent at 15^0^C and much less at higher temperatures). The cloned worms were moved to fresh plates every day till day eight (around the time they stop laying eggs). Worms that escaped, burrowed into the agar, had a bag of worm phenotype or any other morphological defects like bursting of the intestine or protruding vulva were censored and not included in the analysis.

Body bending and pharyngeal pumping analysis was carried out for day-1 adult hermaphrodites. Body bending was analyzed by measuring the number of sinusoidal waves the worm made in one minute (one wave corresponds to one body bend) on a OP50 seeded plate. If the worm stopped moving before one minute, the reading was discarded. For pharyngeal pumping, individual worms were counted for the number of times the pharyngeal contraction took place in 30 seconds.

### Reproductive span and brood size

Reproductive span and brood size were measured by transferring L4 worms onto individual plates. The worms were moved to new plates everyday till they stopped laying eggs. For reproductive span, the number of days till the worm stopped laying eggs was counted. For brood size, the progenies were scored on each plate and the total number was calculated for each worm.

### Stress assays

For the heat stress, day-1 animals were exposed to temperatures from 31^0^C - 41^0^C with an interval of 1^0^C for two hours. Worms were allowed to recover for 24 hours and scored based on movement into three categories: Alive - moving actively and responding to touch, Barely alive - not moving or barely moving, response to touch is not as good as alive worms, and Dead - not moving and not responding to touch. After recovery, it was observed that animals exposed to lower temperatures still had intact bodies but pulsed at higher temperatures had disintegrated. Other *C. briggsae* isolates and mutant cultures were subjected to heat stress at 37^0^C for two hours and scored in the same way.

Osmotic stress was carried out by exposing day-1 worms to hypertonic conditions. NGM agar plates containing varying amounts of sodium chloride (300 and 400mM and 500mM) were made. These plates were seeded with OP50 and stored at 4^0^C for no more than a week. The plates were sealed with parafilm to maintain consistency. 30 - 40 day-1 animals were placed on test plates and scored for survival after 24 hours. Based on the response to touch, the worms were placed into three categories as mentioned in the HS assay.

Oxidative stress was assayed by exposing day-1 worms to 200 mM Paraquat (methyl viologen) solution. The assay was carried out in a 24 well plate format where 35 - 50 day-1 worms were added in two wells for each strain. Worms were screened every hour for survivability up to four hours. Animals that didn’t move in response to touch and had rod-like appearance were counted as dead.

### qRT-PCR

Day-1 AF16 and N2 worms were exposed to HS at 37^0^C for one hour. RNA was collected after a recovery period of 1, 6 and 24 hours. For controls, worms were collected at similar time points without exposure to HS. RNA extraction, DNase treatment and cDNA synthesis was carried out following the protocols mentioned in Ranawade et al. (2018). Levels of the transcription factor *hsf-1* and heat shock chaperon protein (hsp) *hsp-16.2* were determined by qPCR. *pmp-3* and *iscu-1* were used as housekeeping genes for *C. elegans* and *C. briggsae* respectively. The primer pairs used for qPCR are mentioned in Supplementary table 1.

### Statistical analysis

Lifespan graphs and data analysis were carried out using SigmaPlot 14. All other graphs and statistical analyses were performed using GraphPad Prism 9.5.1. RT-qPCR data was analyzed using CFX Maestro 3.1 software (Bio-Rad, Canada; https://www.bio-rad.com/en-ca/product/cfx-maestro-software-for-cfx-real-time-pcr-instruments).

## RESULTS

As part of the systematic characterization of *C. briggsae* as a genetic system, we examined the morphology and other characteristics of the species. Four different wild isolates from different geographical regions (AF16 and QX1410 - tropical, HK104 and VX34 - temperate) were used to investigate and quantify different traits.

### *C. briggsae* is morphologically similar to *C. elegans* but shows subtle differences

The measurements of length and width of *C. elegans* N2 and four *C. briggsae* isolates revealed two facts: one, length is highly variable and, two, *C. briggsae* is slenderer (Figure 1A; Supplementary figure 1A; Supplementary table 2). The VX34 and HK104 isolates were largest and smallest, respectively. The length per unit width of strains ranged from 17.1 to 19.7 with VX34 having the highest value that was significantly different from N2 (*p* < 0.0001) (Figure 1A inset).

**Figure 1:**
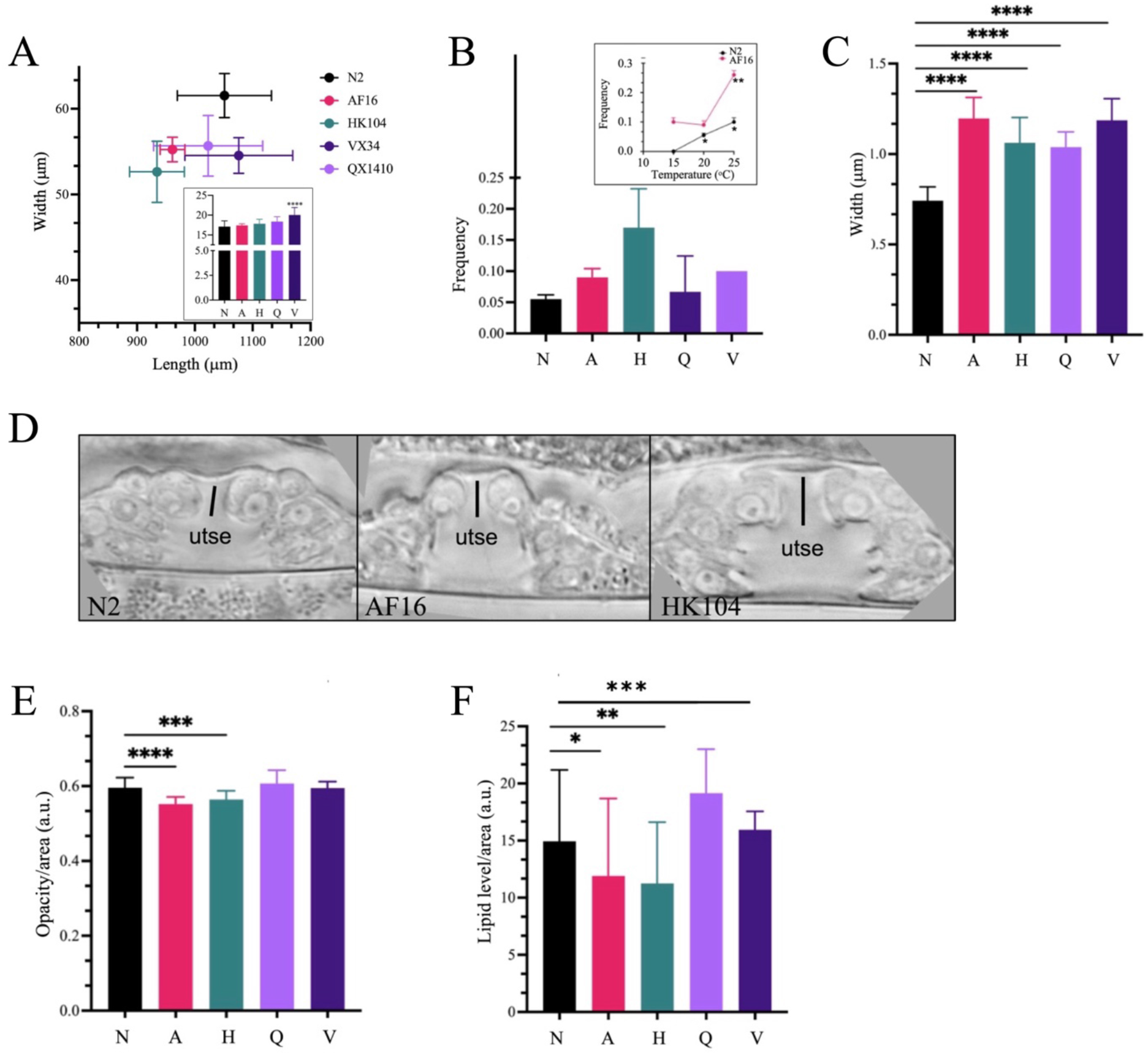
Size, male generation, opacity, and lipid analysis of *C. briggsae* and *C. elegans* isolates. Strain names are abbreviated as N (N2), A (AF16), H (HK104), Q (QX1410), and V (VX34). **A.** Length and width of one day old hermaphrodites. The inset graph shows length to width ratio for each isolate. The N2-VX34 comparison is the only one statistically significant. n = 10 to 17 worms for each strain, combined from two batches. **B.** Male generation frequency at 20^0^C. The inset graph shows frequencies of AF16 and N2 animals at 15^0^C, 20^0^C and 25^0^C. The assay was repeated twice for each strain and temperature condition. **C, D:** Utse thickness of animals. n = 8 - 12 for each strain, combined from two batches. **E:** Opacity (as a measurement of brightness of pixels) of one day old hermaphrodites. The assay was carried out in triplicates with n = 20 to 30 for each strain. **F:** Lipid staining using Oil Red O technique for AF16, HK104 and N2 day-1 animals. The assay was carried out in triplicates with n = 20 to 30 for each strain. **A-C, E, F.** Data is represented as mean ± SD. One-way ANOVA with Dunnett’s multiple comparisons test was used to analyze data. Statistically significant values are indicated by star (*): * *p* < 0.05; ** *p* < 0.01; *p* < 0.001; **** *p* < 0.0001; ns, not significant.

Casual observations of AF16 and N2 animal tracks on solid agar culture plates hinted at some differences in movement characteristics. To examine this further, we quantified the sinusoidal wave pattern and found that AF16 day-1 adults had a comparatively shorter amplitude (Supplementary figure 2A). However, the amplitude per unit length of both species was comparable, suggesting that both species have a similar sinusoidal wave pattern of movement (Supplementary figure 2B).

Nigon and Dougherty had reported earlier that *C. elegans* and *C. briggsae* produce more males spontaneously at higher temperatures due to meiotic non-disjunction of sex chromosome (Nigon and Dougherty, 1949). During routine culturing of *C. briggsae* AF16, we observed a higher incidence of males compared to *C. elegans* N2. To follow up on this, males were quantified in the F1 progeny of unmated hermaphrodites of N2 and *C. briggsae* isolates. At 20^0^C all strains showed similar frequency of males that ranged from 0.05 to 0.1% (Figure 1B). The HK104 showed a larger variation but was not significantly different other cultures. The N2 and AF16 animals were examined at two additional temperatures, 15^0^C and 25^0^C. The results revealed roughly 2-3-fold more males in AF16 compared to N2 (Figure 1B inset). The frequency was much higher at 25^0^C, likely due to increased non-disjunction. No N2 males were recovered at 15^0^C. It is worth mentioning that males in *C. briggsae* AF16 cultures persisted even after brief starvation and could be propagated by chunking.

It was previously observed that the uterine-seam cell syncytium (utse) at the vulval apex in L4 stage animals is slightly thicker in *C. briggsae* AF16 compared to *C. elegans* N2 (Gupta and Sternberg, 2003). We expanded on this observation by examining other *C. briggsae* isolates and found the utse to be ~1.5x thicker in each strain when compared to N2 (Figure 1C, D). It should be mentioned that in despite differences in the utse morphology, the egg laying responses of two species appeared to be similar.

Among other observations, it was noted that *C. briggsae* AF16 animals are comparatively more transparent than *C. elegans* N2. Measurements of opacity confirmed that it was indeed the case. Thus, *C. elegans* N2 was significantly darker than *C. briggsae* AF16 and HK104 animals (Figure 1E). However, opacities of VX34 and QX1410 isolates were comparable to *C. elegans* N2, suggesting variability in the trait. To examine if the opacity would correlate with lipid levels, we performed Oil Red O staining. The results did not show a clear trend (Figure 1F). Further investigations of the relationship between lipids and transparency involved making use of *C. elegans* mutants, *daf-2 (e1370)* and *pry-1 (gk3682)*, that have high and low lipid contents, respectively (O’Rourke et al., 2009; Ranawade et al., 2018). The data showed that both mutant stains are opaquer than N2 (Supplementary figure 3), leading us to conclude that while lipids may affect opacity, other factors also contribute to differences in body color.

### *C. briggsae* grows, survives, and reproduces at higher temperature better than *C. elegans*

In addition to the above visual characteristics, we examined physiological features of *C. briggsae* and compared those with *C. elegans*. Both nematodes thrive in organic rich matter namely compost and rotting fruits. (Kiontke and Sudhaus, 2006). Even though they are found in similar environments, there are differences in their seasonal abundances with *C. briggsae* preferring higher temperature conditions. (Félix and Duveau, 2012). To investigate the temperature trait further, we looked at various characteristics of N2 and AF16 strains grown at different temperatures.

The examination of time to reach adulthood revealed striking differences wherein at 15^0^C *C. briggsae* AF16 took 15.8% longer time than *C. elegans* to reach sexual maturity (112.70 +/− 1.15 hours and 97.33 +/− 1.15 hours respectively; *p* < 0.0001) but 16.8% shorter time at 25^0^C (42.47 +/− 1.36 hours and 51.33 +/− 1.15 hours respectively; *p* < 0.01) (Figure 2A). At 20^0^C, growth rates did not show a significant difference (67.33 +/− 2.51 hours for *C. briggsae* and 70.33 +/− 1.52 hours for *C. elegans*; Figure 2A). The differences in developmental timings were also evident in temporal analysis of animals reaching adulthood (Figures 2B, C, D).

**Figure 2:**
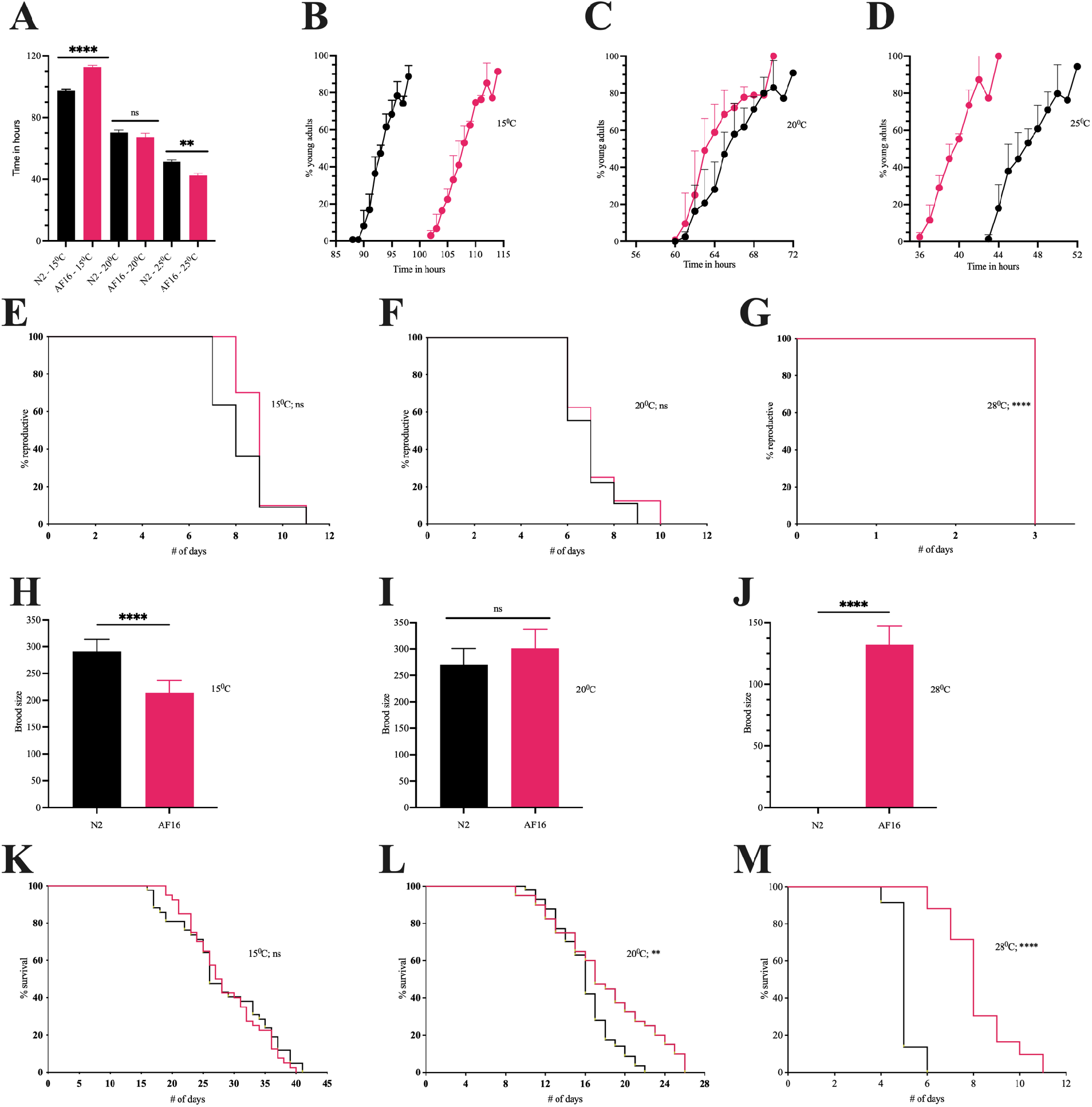
Developmental time, brood size, and life span of AF16 and N2 at various temperatures. **A.** Time taken by 80% of AF16 and N2 population to reach young adults at 15^0^C, 20^0^C and 25^0^C. n = 109 to 159 in a total of three batches for both species and each temperature condition. **B-D.** Temporal analysis at 15^0^C (B), 20^0^C (C) and 25^0^C (D). **E-G.** Reproductive spans of AF16 and N2 at 15^0^C (E), 20^0^C (F), and 28^0^C (G). **H-J.** Brood size of AF16 and N2 hermaphrodites at 15^0^C (H), 20^0^C (I), and 28^0^C (J). For **E-J,** 8-10 worms were measured for each temperature and each strain in a total of two batches. **K-M.** Life span of AF16 and N2 at 15^0^C (K), 20^0^C (L), and 28^0^C (M). n = 2 batches of pooled worms, with minimum 20 worms scored in each batch for each strain and temperature condition. **A-D and H-J**: Data is represented as mean ± SD. Statistical tests used: **A, H-J:** unpaired t test. **E-G, K-M:** Kaplan-Meier Log Rank (Mantel-Cox) test. Statistically significant values are indicated by asterisk (*): * *p* < 0.05; ** *p* < 0.01; **** *p* < 0.0001; ns, not significant.

Similar to the growth rates, brood size and reproductive span of animals were also affected by temperature. Specifically, the brood size of N2 was higher than that of AF16 at 15^0^C but comparable at 20^0^C. At 28^0^C, N2 was sterile whereas AF16 produced progeny (Figures 2E, F, G, H, I, J). The mean reproductive spans are listed in Table 1, which show an inverse relationship with temperature. It is interesting to note that while the brood size of AF16 was significantly lower than N2 at 15^0^C, the reproductive span was comparable at this temperature (Figures 2E, H and Table 1).

**Table 1.** Reproductive span of AF16 and N2 animals at different temperatures. Each assay was conducted in two batches. n represents the total number of animals from combined batches. NA, Not applicable (animals are sterile). See figure legends for details of statistical analyses.

| Genotype | Temperature | Mean lifespan ± standard error | Median lifespan | Reproductive span | n | *p* value |
| --- | --- | --- | --- | --- | --- | --- |
| N2 | 15°C | 8.18 ± 0.37 | 8 | 11 | 11 | 0.233 |
| AF16 | 15°C | 8.80 ± 0.29 | 9 | 11 | 10 | |
| N2 | 20°C | 6.89 ± 0.35 | 7 | 9 | 9 | 0.604 |
| AF16 | 20°C | 7.13 ± 0.48 | 7 | 10 | 8 | |
| N2 | 28°C | NA | NA | NA | NA | <0.001 |
| AF16 | 28°C | 3 ± 0.00 | 3 | 3 | 8 | |

We also examined the survivability of nematodes by growing them at different temperatures. The data is presented in Figures 2K, L, M and Table 2. Earlier, *C. briggsae* AF16 animals were reported to live longer than *C. elegans* N2 at 20^0^C (Mallick et al., 2020). We observed similar results. Despite differences in the lifespan, the physiological markers of aging, i.e., pharyngeal pumping and body bending frequencies, were similar in both species (Figure 2L, Supplementary figure 4, and Table 2). Lifespan studies were also carried out at two additional temperatures, one at a lower end (15^0^C) and the other at a higher end (28^0^C). The data showed that *C. briggsae* has a significantly longer lifespan than *C. elegans* at 28^0^C but comparable at 15 ^0^C (Figures 2K, M; Table 2). Taken together, these results show that aging characteristics of *C. elegans* N2 and *C. briggsae* AF16 differ from growth rate and brood size traits when reared at different temperatures.

**Table 2.** Lifespan analysis of AF16 and N2 animals grown at different temperatures. Each assay was conducted in two batches. ‘n’ represents the total number of animals from combined batches. See figure legends for details of statistical analyses.

| Genotype | Temperature | Mean lifespan ± standard error | Median lifespan | Maximum life span | n | *p* value |
| --- | --- | --- | --- | --- | --- | --- |
| N2 | 15<sup>0</sup>C | 28.31 ± 1.16 | 26 | 41 | 42 | 0.161 |
| AF16 | 15<sup>0</sup>C | 27.4 ± 0.92 | 26 | 40 | 40 | |
| N2 | 20<sup>0</sup>C | 16.03 ± 0.39 | 16 | 22 | 40 | 0.002 |
| AF16 | 20<sup>0</sup>C | 17.97 ± 0.80 | 17 | 26 | 57 | |
| N2 | 28<sup>0</sup>C | 5.05 ± 0.04 | 5 | 6 | 92 | <0.001 |
| AF16 | 28<sup>0</sup>C | 8.16 ± 0.14 | 8 | 11 | 118 | |

### *C. briggsae* has higher thermal resistance than *C. elegans* but is comparatively more sensitive to other forms of stress

The experiments described above show that *C. briggsae* is better adapted to higher temperatures. To investigate this further, we examined the resistance of *C. briggsae* to acute heat treatments. For this N2 and AF16 were exposed to temperatures ranging from 32^0^C to 41^0^C for two hours. Examination of animals after heat pulses showed that while some were moving actively, others were barely mobile, or appeared dead (Figure 3A). It was observed that AF16 is more tolerant to higher temperatures than N2 and can survive heat shocks up to 39^0^C (Figure 3A). At 37^0^C, there is a clear difference between the two species; while AF16 is mostly alive (>90%), more than 65% of N2 animals are either barely alive or dead. We tested three other *C. briggsae* isolates (HK104, VX34 and QX1410) and observed a similar heat resistant response (Figure 3B). There was also a clear difference in the movement characteristics of two species following heat treatment with *C. briggsae* appearing more active and healthier than *C. elegans* (Figures 3C, D).

**Figure 3:**
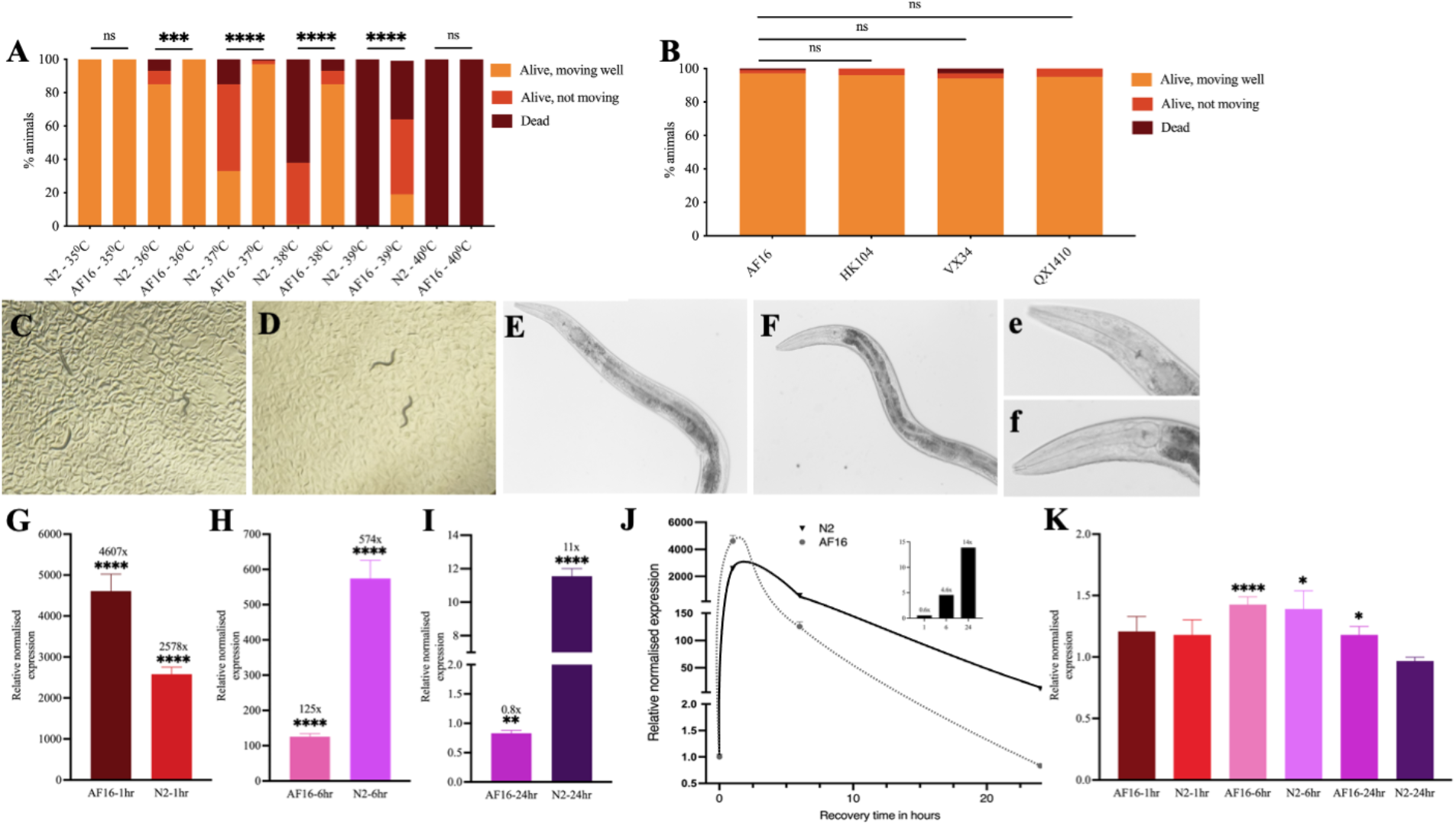
Effect of different heat stress conditions on AF16 and N2. **A.** Day-1 wild type *C. elegans* (N2) and *C. briggsae* (AF16) adults were exposed to a range of temperatures for 2 hours for heat stress. **B.** *C. briggsae* isolates HK104, VX34 and QX1410 were exposed to heat stress at 37C for 2 hours. For **A**, **B**, the assay was repeated at least twice for each strain and temperature condition (n = 30 to 40 worms in each batch). **C.** Day-1 N2 animals after a recovery period of 24 hours following heat shock at 37C for 2 hours. **D.** Day-1 AF16 animals after a recovery period of 24 hours following heat shock at 37C for 2 hours. **E.** N2 day-1 adult **F.** AF16 day-1 adult. The 20x magnified images of E and F are shown in **e** and **f**, respectively. **G-I.** Relative normalized expression of *hsp-16.2* in heat shocked worms of AF16 and N2 after a recovery period of 1, 6 and 24 hours, respectively. **J.** the graph shows fold change in expression of *hsp-16.2* for AF16 and N2. The inset shows ratios of relative expression at each recovery time point. **K.** Relative normalized expression of *hsf-1* in heat shocked worms of AF16 and N2 after a recovery period of 1, 6 and 24 hours. For **G-I, K**, the qPCR analysis was repeated three times for each recovery period and each strain. Data is represented as mean ± SEM. Statistical tests used: **A**, **B.** Chi-square test. **G-I, K:** t-test. Statistically significant values are indicated by asterisk (*): * *p* < 0.05; ** *p* < 0.01; ns, not significant.

We also examined treated animals under the Nomarski microscope, which showed differences in internal organs. For example, pharyngeal bulb and intestinal structures had less well-defined outlines (Figures 3E, F and insets); although the basis of such changes, e.g., due to tissue damage or altered membrane composition, remains to be investigated.

Since the heat shock response is known to activate expression of a battery of chaperons to protect proteins from misfolding (Jovic et al. 2017), we investigated whether expression dynamics of two major factors, the heat shock factor 1 (*hsf-1*) and the cytosolic heat shock protein 16.2 (*hsp-16.2*), would correlate with heat tolerance of the two species. To this end, animals were subjected to 1 hour heat shock at 37^0^C, following which *hsf-1* and *hsp-16.2* levels were measured at different time points. The results showed differences in *hsp-16.2* activation. Whereas in *C. elegans*, the fold changes were roughly 2,500x, 500x, and 10x after recovery periods of 1 hr, 6 hr, and 24 hr; respectively, the corresponding increases in *C. briggsae* were 4600x, 125x, and 1x (Figures 3G, H, I). These results show that *hsp-16.2* is robustly activated in both species following the heat shock. Furthermore, the levels are almost twice as high in *C. briggsae* compared to *C. elegans* at 1 hr time point; and show a faster decline in *C. briggsae* (Figure 3J, inset). Unlike *hsp-16.2*, *hsf-1* transcription was not greatly affected, i.e., no change after 1 hr, less than 1.5x increase after 6 hrs, and very little change after 24 hrs (Figure 3K). These results show that while heat shock genes were activated in both species, *hsp-16.2* exhibits a greater transcriptional change in *C. briggsae* following heat stress.

In addition to *hsp-16.2* and *hsf-1*, we looked for other factors that may contribute to heat resistance of *C. briggsae*. This was done using a collection of mutants isolated earlier from forward genetic screens in our lab and other labs (Gupta et al., in preparation; www.briggsae.org). Specifically, 15 strains exhibiting a range of phenotypes (Unc, Sma, Bli, Dpy, and Lin) were selected. The animals were subjected to a 2 hr heat shock at 37^0^C and survival was measured after 24 hr of recovery (Supplementary Figure 5). The *Cbr-lin-11(sy5336)* adults showed sensitivity to heat stress that was attributed to their egg-laying defective (Egl) phenotype since L4 worms were comparable to controls. A similar result was obtained for *Cbr-unc(sy5010)*, although in this case L4 animals were weakly sensitive. The *Cbr-unc(sy5077)* adults showed high sensitivity to heat stress, most likely due to animals being severely uncoordinated and unhealthy (data not shown). Overall, we did not find any obvious candidate affecting the heat resistance of animals. Since most of the mutants used in this study remain uncharacterized, a firm conclusion about the role of these genes, including any potential genetic redundancies, cannot be made.

In addition to heat, we also examined the responses of *C. briggsae* and *C. elegans* to two other stresses, i.e., oxidative stress and osmotic stress. Interestingly, *C. briggsae* showed increased sensitivity to both these chemicals. Thus, after 4 hours of paraquat exposure (for oxidative stress), fewer AF16 and HK104 animals were alive compared to N2 (Figure 4A). A similar trend was observed following 24 hours of NaCl exposure (for osmotic stress) (Figure 4B). Thus, *C. briggsae* is not resistant to all forms of stress, suggesting that higher thermal tolerance is a specific response of this species.

**Figure 4:**
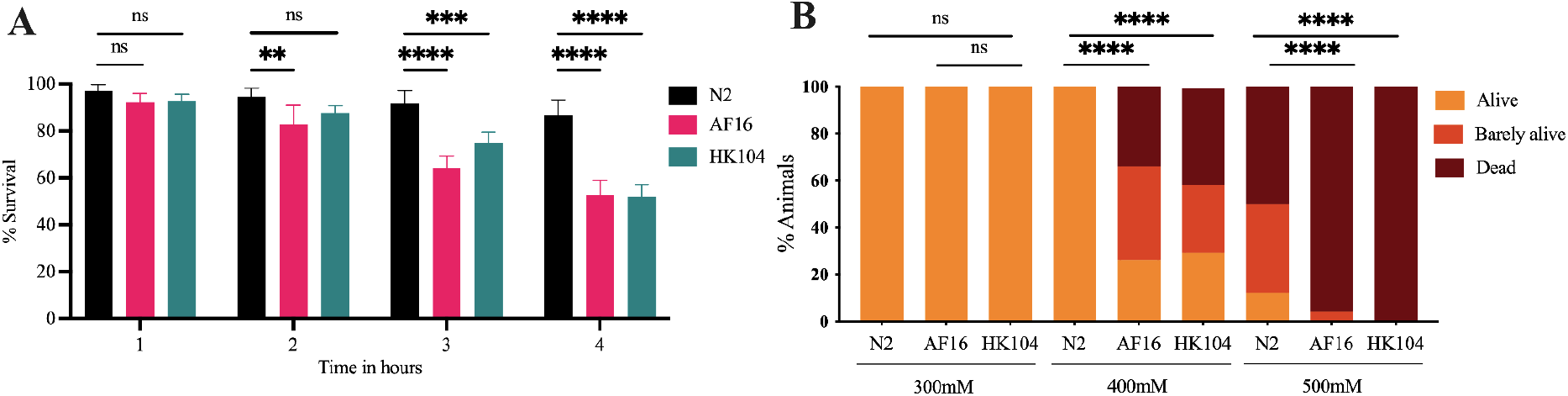
Effect of oxidative and osmotic stress on AF16, H104 and N2. **A.** Day-1 worms were treated with 200mM PQ for 4 hours. Survival was measured after each hour. n = 112 - 141 in a total of 2 batches (2 wells per batch) for all strains. **B.** Day-1 AF16, HK104 and N2 animals were exposed to 300mM, 400mM and 500mM NaCl and survivability was measured after 24 hours. n = 37 to 78 in a total of 2 batches for all strains. **A.** Data is represented as mean ± SD. Statistical tests used: **A.** Chi-square test. **B.** Two-way ANOVA with Tukey’s post hoc test. Statistically significant values are indicated by asterisk (*). * *p* < 0.05; ** *p* < 0.01; *** *p* < 0.001; **** *p* < 0.0001; ns, not significant.

## DISCUSSION

This study focused on morphological and physiological characteristics of *C. briggsae* temperate and tropical isolates and their comparisons with the established *C. elegans* wild-type laboratory strain N2. Our results extend the findings on similarities and differences between the two species. Specifically, we found that while *C. briggsae* isolates were comparatively thinner than *C. elegans* N2, the animals exhibited greater variability in length, similar to that reported for *C. elegans* wild isolates (McCulloch and Gems, 2003). Variations in size have also been reported for other nematodes. *C. inopinata* adults are roughly 1.5x longer than *C. elegans* however, interestingly, their dauers are comparatively smaller (Hammerschmith et al., 2022). Another study by Flemming et al. (2000) showed that the length of Rhabditida species varies by six fold. Studies in *C. elegans* have shown that body size is affected by both genetic and environmental factors (e.g., see Van Voorhies, 1996; Kammenga et al., 2007; Gumienny and Savage-Dunn, 2013; Maulana et al., 2022). Genetic factors include genes such as the sma (small) class that belong to the TGFβ signaling (Gumienny and Savage-Dunn, 2013) and *tra-3/Calpain 5* (Kammenga et al., 2007).

Our study revealed differences in the utse morphology between the two species. In the L4 stage larvae, utse appears as a thin membrane (hymen) at the vulva apex. We found the membrane to be 1.5x thicker in *C. briggsae* isolates compared to *C. elegans* N2. Whether such a difference has an impact on egg-laying characteristics of these species remains to be seen.

We also observed changes in the opacity of *C. briggsae* isolates and investigated its relationship to lipid content. Lipid measurements in wild strains and *C. elegans* mutants *daf-2* and *pry-1* with high and low lipids, respectively, suggested that lipids are not the sole determinants of opacity, raising the possibility that factors such as cuticle composition may also be involved (Supplementary figure 3).

Among other characteristics, we found a higher frequency of spontaneous males (2 - 3 folds) in AF16 compared to N2. Considering that *C. elegans* wild strains exhibit significant differences in the proportion of males (Anderson et al., 2010), these data suggest that non-disjunction frequency varies greatly among species and isolates. Whether such differences result from altered expression of any of the known genes linked to male production (Hodgkin et al., 1979; Lim et al., 2021) remains to be investigated. A related observation was that unlike *C. elegans* N2 where the frequency of males rapidly declines in actively growing cultures, *C. briggsae* males persist even after starvation and repeated transfers of cultures. This may be due to *C. briggsae* males being more efficient in mating. In agreement with this, (Garcia et al., 2007) had shown earlier that *C. briggsae* males are better at mating with young, 1-day old adult hermaphrodites than *C. elegans* males.

Finally, we quantified the sinusoidal movement of animals on culture plates, which revealed that while the absolute amplitude of AF16 is comparatively shorter, the amplitude per unit length is similar to N2 (Supplementary figure 2).

The above results add to the existing body of work on morphological features of *C. briggsae* and their differences from *C. elegans*. These include anterior placement of excretory duct (Wang and Chamberlin, 2002) altered arrangements of bursal rays in male tail (Fitch, David H.A. Emmons, 1995; Fitch, 1997) reduced competence of P3.p vulval precursor cell (Delattre and Felix, 2001), lack of systemic RNAi (Winston et al., 2007), and susceptibility towards viral infections (Félix et al., 2011; Franz et al., 2012; Frézal et al., 2019). Experiments involving physiological responses have also uncovered some changes in *C. briggsae*, such as the faster electrotaxis speed in a microfluidic channel (Rezai et al., 2011) and lack of dauers at high temperature (Inoue et al., 2007). Other non-physical differences include alterations in signaling pathways (Félix, 2007; Hoyos et al., 2011; Mahalak et al., 2017) and telomeric small RNAs (Frenk et al., 2019).

A major finding of our work is that *C. briggsae* grows, survives, and reproduces better than *C. elegans* at higher temperatures. We found that growth rates of AF16 animals to be faster at 25^0^C compared to N2 but slower at 15^0^C (roughly 16% - time difference for each). A similar trend was observed for the brood size, suggesting that AF16 is better adapted to a warmer climate. Consistent with these data, a previous work had reported the fecundity of *C. briggsae* isolates at different temperatures and showed that both temperate and tropical isolates reproduced robustly when grown between 20^0^C and 28^0^C (Prasad et al., 2011). Studies in *C. elegans* have shown the effect of temperature on brood size. McMullen et al. (2012) examined the egg laying of N2 animals and showed that at 28^0^C, the average number of eggs laid was reduced significantly whereas no eggs were laid at 30^0^C. This phenomenon is not unique to N2, as other *C. elegans* isolates exhibited similar characteristics (Petrella, 2014). Another study by Poullet et al. (2015) reported intra and inter species thermal plasticity in spermatogenesis, oogenesis, mitosis, and meiosis. These findings together with data presented here advance our understanding of the effect of temperature on growth, reproduction, and brood size of *Caenorhabditis* species.

Lifespan of *C. elegans* and *C. briggsae* isolates from different geographical locations has been reported (Gems and Riddle, 2000; Joyner-Matos et al., 2009; Yujin et al., 2016). It is interesting to note that a correlation between lifespan and region of isolation was observed for *C. briggsae* but not for *C. elegans* (Joyner-Matos et al., 2009; Yujin et al., 2016). Specifically, *C. briggsae* strains from temperate regions were shown to have a longer lifespan than those from equatorial and tropical regions. Our analysis reveals that while *C. briggsae* AF16 has a longer lifespan at 20^0^C and 28^0^C, it is comparable to N2 at 15^0^C. To further investigate the thermal responses of *C. briggsae*, we performed acute treatments at high temperatures. The results revealed that AF16 and other isolates are more resistant to heat stress than *C. elegans* N2. Interestingly, *C. briggsae* showed greater sensitivities to oxidative and osmotic stress, suggesting the resistance to heat stress is a specific characteristic of this species. While the underlying basis of differences in stress responses between the two species is currently unknown, factors such as cuticle composition, lipids, and heat shock chaperons may be involved. A recent study showed the role of permeability-determining (PD) collagens in maintaining cuticle structure, such that their loss leads to enhanced sensitivity to paraquat, levamisole and ivermectin (Sandhu et al., 2021). We have observed significant differences in the expression of collagen genes between *C. briggsae* and *C. elegans* adults (van den Berg et al., in preparation). Other cellular components could play roles as well, e.g., Lamitina et al. (2006) reported the involvement of glycerol phosphate dehydrogenases in protecting *C. elegans* against osmotic stress. Thus, expression differences in these and other genes between the two species may contribute to variations in responses to temperature stress.

Lipids and fatty acids are known to affect sensitivity of animals to environmental stress (see review by Los and Murata, 2004). Horikawa and Sakamoto (2009) reported earlier that fatty acids regulate heat, oxidative and osmotic stress in *C. elegans*. Specifically, when animals were exposed to high temperatures, a reversible change in the fatty acid composition of the plasma membrane was observed. This change involved an increase in saturated fatty acids and a decrease in unsaturated fatty acids (Horikawa and Sakamoto, 2009; Chauve et al., 2021), thus conferring resistance to high temperatures. In an opposing trend, unsaturated fatty acids were shown to play a role in oxidative stress response, such that a decrease in their levels causes sensitivity to oxidative stress. It is conceivable that the proportion of saturated to unsaturated fatty acid may differ between the two species, thereby causing changes in their stress tolerance. Future studies on fatty acid composition are needed to investigate this possibility.

In addition to lipid and cuticle compositions, heat shock proteins such as the transcription factor HSF-1 and various chaperons (HSPs) are also differentially regulated to affect stress responses of animals (Higuchi-Sanabria et al., 2018). Experiments have shown that exposure to a variety of stresses cause significant changes in the expression and subcellular localization of heat shock proteins thereby enabling animals to endure the stressful conditions (Labbadia and Morimoto, 2015). The levels of *hsp-70* and *hsp-16.11* increase after exposure to HS and gradually decline to normal levels by 8 hours into recovery (Labbadia and Morimoto, 2015). Our findings align with the above data showing *hsp-16.2* levels rising initially to more than two thousand folds in *C. elegans* (2,579x) and about four and a half thousand in *C. briggsae* (4,608x) following exposure to heat stress that gradually declined afterwards. It is conceivable that a greater increase in *hsp-16.2* may confer better protection on *C. briggsae*. In this regard, it is worth mentioning that *hsp-16.2* levels in *C. elegans* are reported to correlate with beneficial effects, i.e., more *hsp-16.2* means better protection against stress and increased longevity of animals (Walker et al., 2001; Cypser et al., 2013; Mendenhall et al., 2017; Burnaevskiy et al., 2019).

Interestingly, while *C. briggsae* showed a much higher increase in *hsp-16.2* expression, it also exhibited a steeper decline with levels going back to normal after 24 hours. In contrast, expression dynamics in *C. elegans* were much shallower and levels remained significantly high after 24 hours (~10x). It remains to be investigated whether the above differences in *hsp-16.2* expression are intrinsic characteristics of these two *Caenorhabditis* species, i.e., chaperon expression levels and dynamics in *C. elegans* following heat treatment will differ from *C. briggsae* regardless of the temperature tested, or be dependent on the condition of the assay, i.e., expression trends in both species will be similar at lower temperatures that don’t affect *C. elegans* viability. Unlike *hsp-16.2*, the levels of heat-shock transcription factor, *hsf-1*, were not greatly affected. This is not unexpected since *hsf-1* is mostly regulated post-translationally following heat stress (Higuchi-Sanabria et al., 2018). It was shown that HSF-1 is nuclear localized, leading to gene expression changes to confer protection against heat stress (Morton and Lamitina, 2013).

Our findings of differences in the acute heat shock response between *C. briggsae* and *C. elegans* is significant. We found that both temperate and tropical isolates of *C. briggsae* show similar resistance to heat stress, suggesting that a higher heat tolerance is an inherent trait of this species. Interestingly, *C. briggsae* showed greater sensitivities to osmotic and oxidative stress. These findings indicate changes in gene function and pathways in these two species, which confer resistance to specific forms of stress. Further characterization of thermal tolerance in *C. briggsae* promises to uncover mechanisms that could help understand how cells and tissues respond to heat stress across metazoans.

## ACKNOWLEDGEMENTS

We sincerely thank many colleagues for ideas and thoughtful comments on various experiments described in this manuscript. We also thank Erik Anderson for providing the VX34 and QX1410 isolates. Several Gupta lab members assisted with techniques and reagents. Some of the morphological studies originated from early discussions with Takao Inoue. Funding for this work was provided by the Natural Science and Engineering Research Council of Canada Discovery grant to BPG.

## Supplementary Figures

**Supplementary figure 1.**
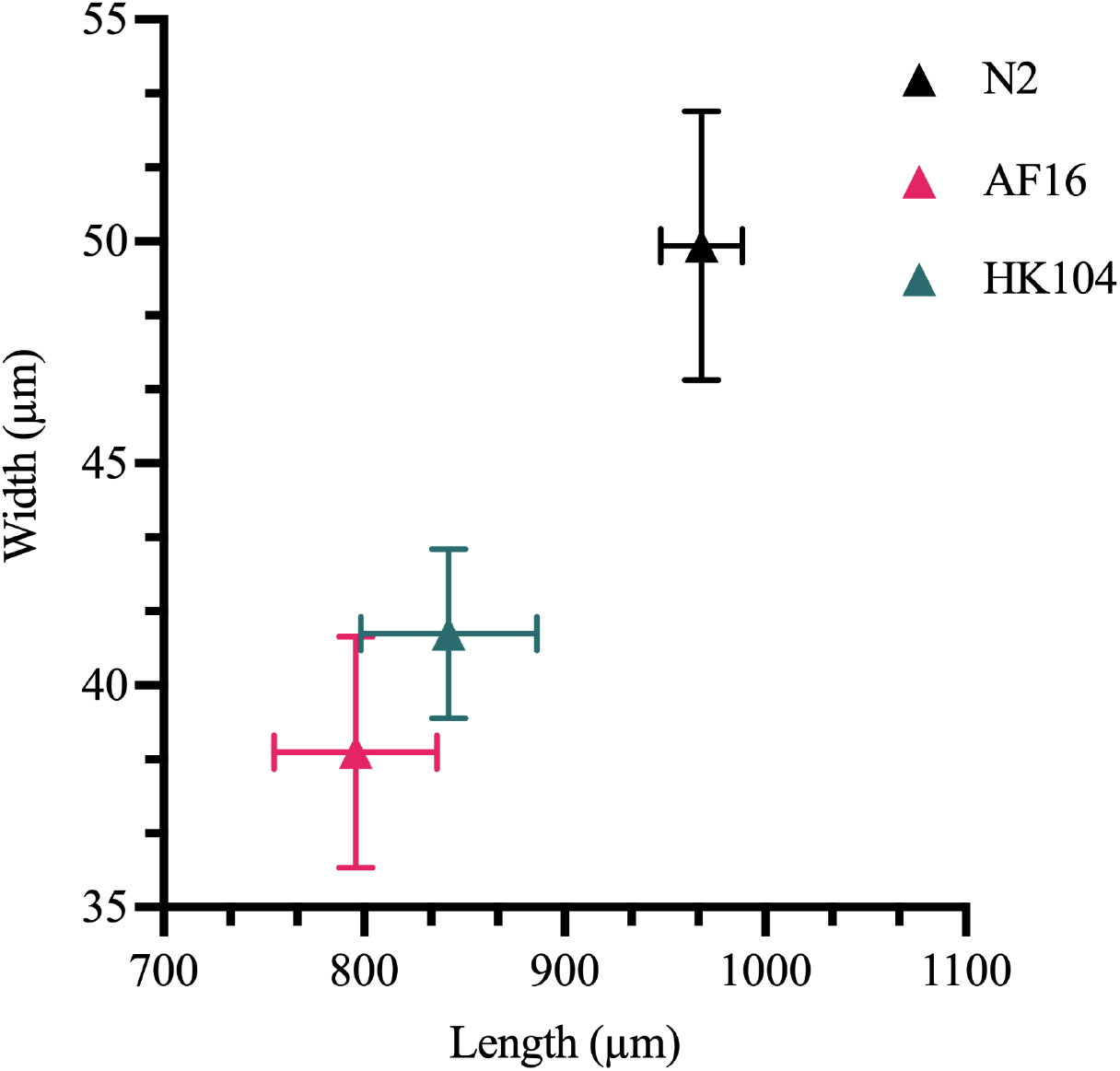
Measurements of day-1 adults of AF16, HK104 and N2 males. Length and width of males are plotted. n = 10 animals for each strain, combined from two batches. Data is represented as mean ± standard deviation. All measurements are in micrometers. One-way ANOVA using Dunnett’s multiple comparison test was used to analyze data. The differences for length and width between N2 and each of the *C. briggsae* isolates were calculated and found to be significant (*p* < 0.0001 in each case).

**Supplementary figure 2.**
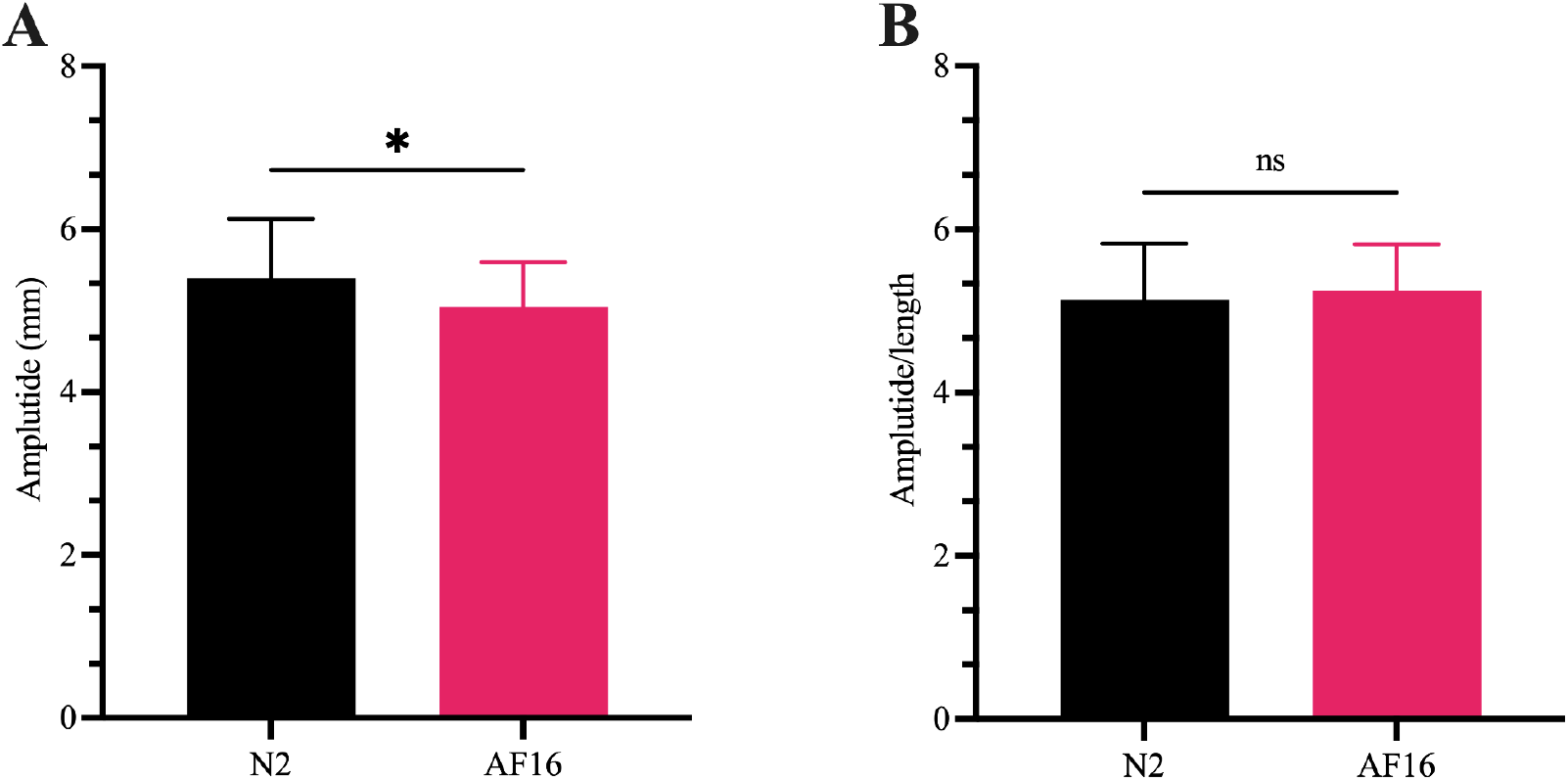
Movement analysis of AF16 and N2. **A.** Amplitude of AF16 and N2 day-1 adults was measured. AF16 has a shorter amplitude compared to N2. **B.** There is no significant difference in the amplitude per unit length between AF16 and N2. Four worms (10 waves per worm) were measured for each strain in a total of two batches. **A, B.** Data is represented as mean ± SD. Statistical analysis was done using the student’s unpaired *t* test. Significant values are indicated by asterisk (*). *: (*p* < 0.05). ns, not significant.

**Supplementary figure 3.**
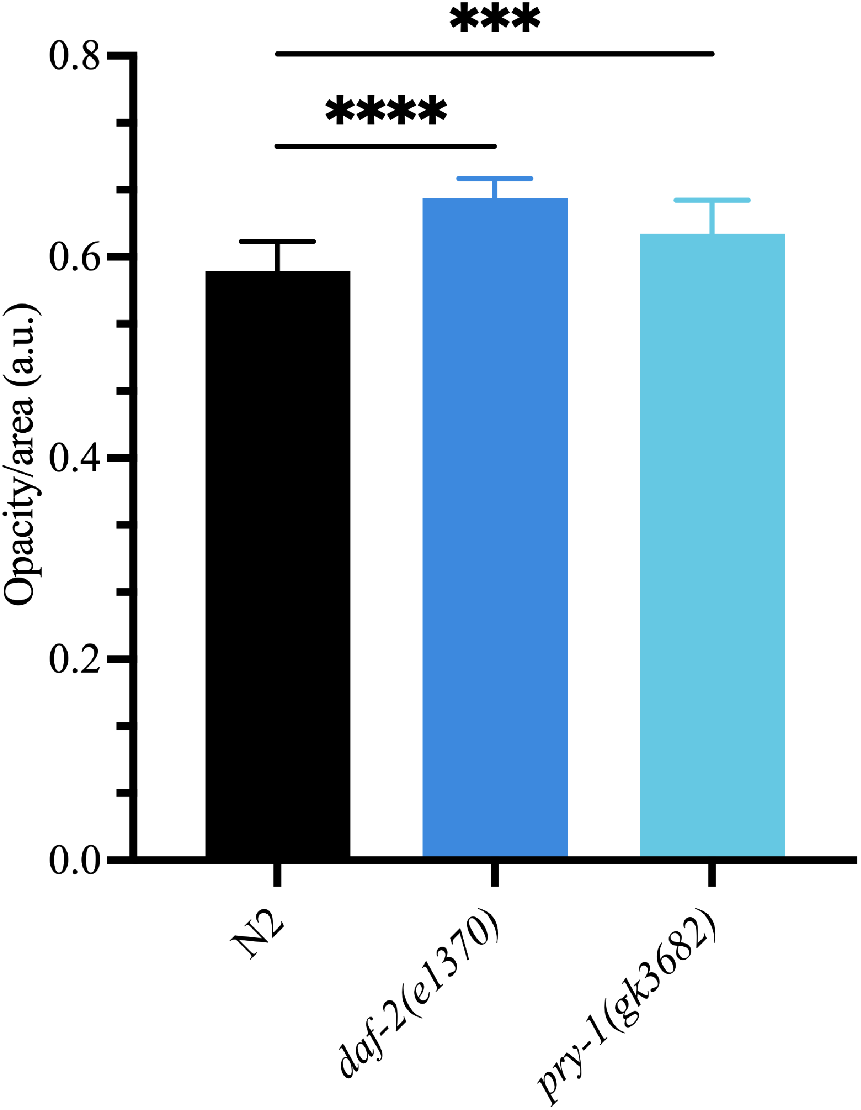
Opacity of N2, *daf-2(e1370)* and *pry-1(gk3682)*. Day-1 animals of N2, *daf-2(e1370)* and *pry-1(gk3682)* were measured for opacity. One-way ANOVA using Dunnett’s multiple comparison test shows significant differences between N2 and *daf-(e1370)*, and N2 and *pry-1(gk3682)*. n = 17 to 27 animals in a total of 3 batches for all strains. Data is represented as mean ± SD. Statistically significant values are indicated by asterisk (*). *** (*p* < 0.001); **** (*p* < 0.0001).

**Supplementary figure 4.**
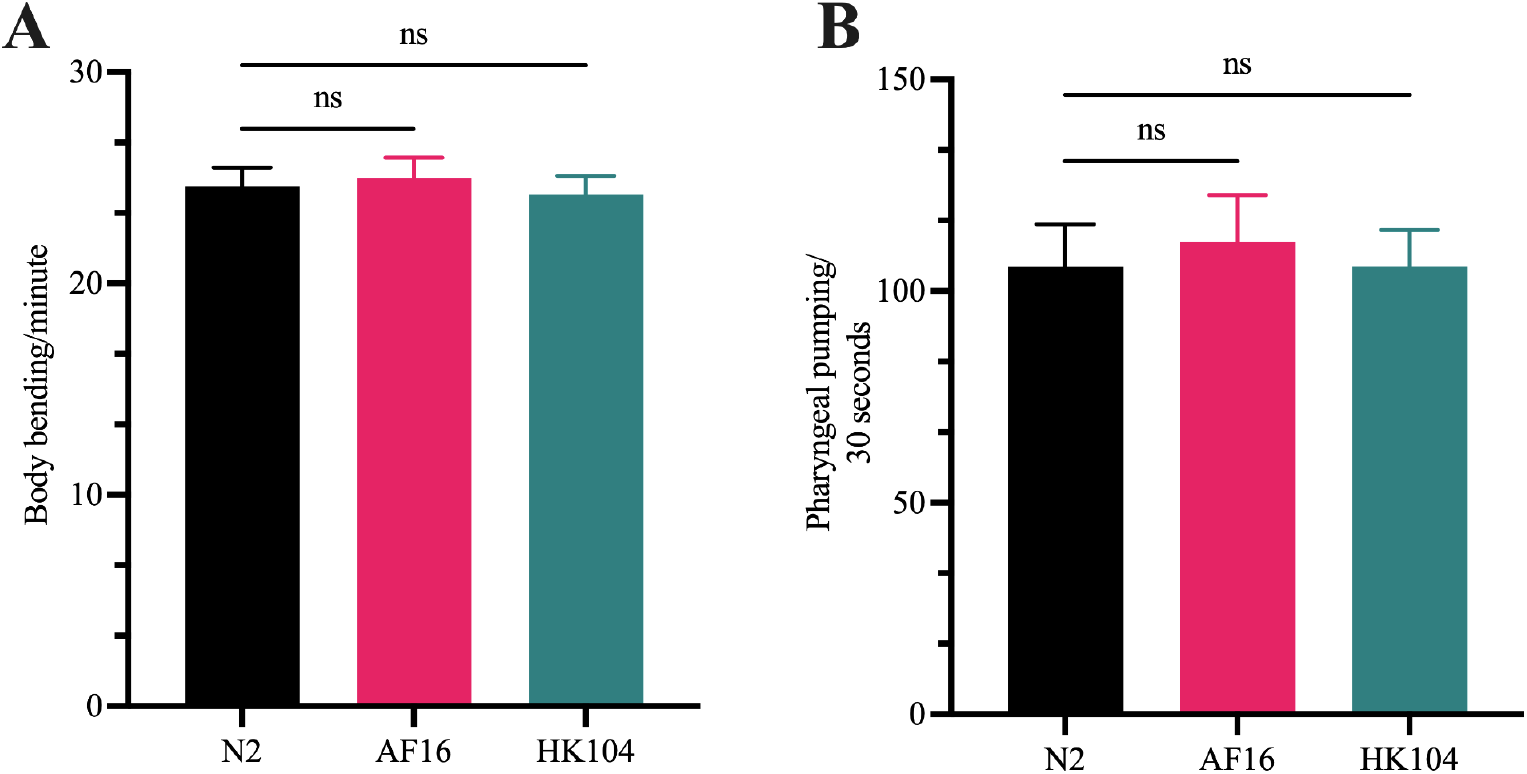
Body bending and pharyngeal pumping of AF16, HK104 and N2 at 20 ^0^C. **A.** Day-1 animals of N2, AF16 and HK104 were analyzed for their body bends per minute. There is no significant difference between N2 and both C. *briggsae* strains. **B.** Pharyngeal pumping of AF16, HK104 and N2 revealed no significant difference (ns) between N2 and AF16, and N2 and HK104. n = 20 - 30 animals in a total of 3 batches for all strains. **A, B.** Data is represented as mean ± SD. Statistical analysis was carried out using one-way ANOVA using Dunnett’s multiple comparison test. ns, not significant.

**Supplementary figure 5.**
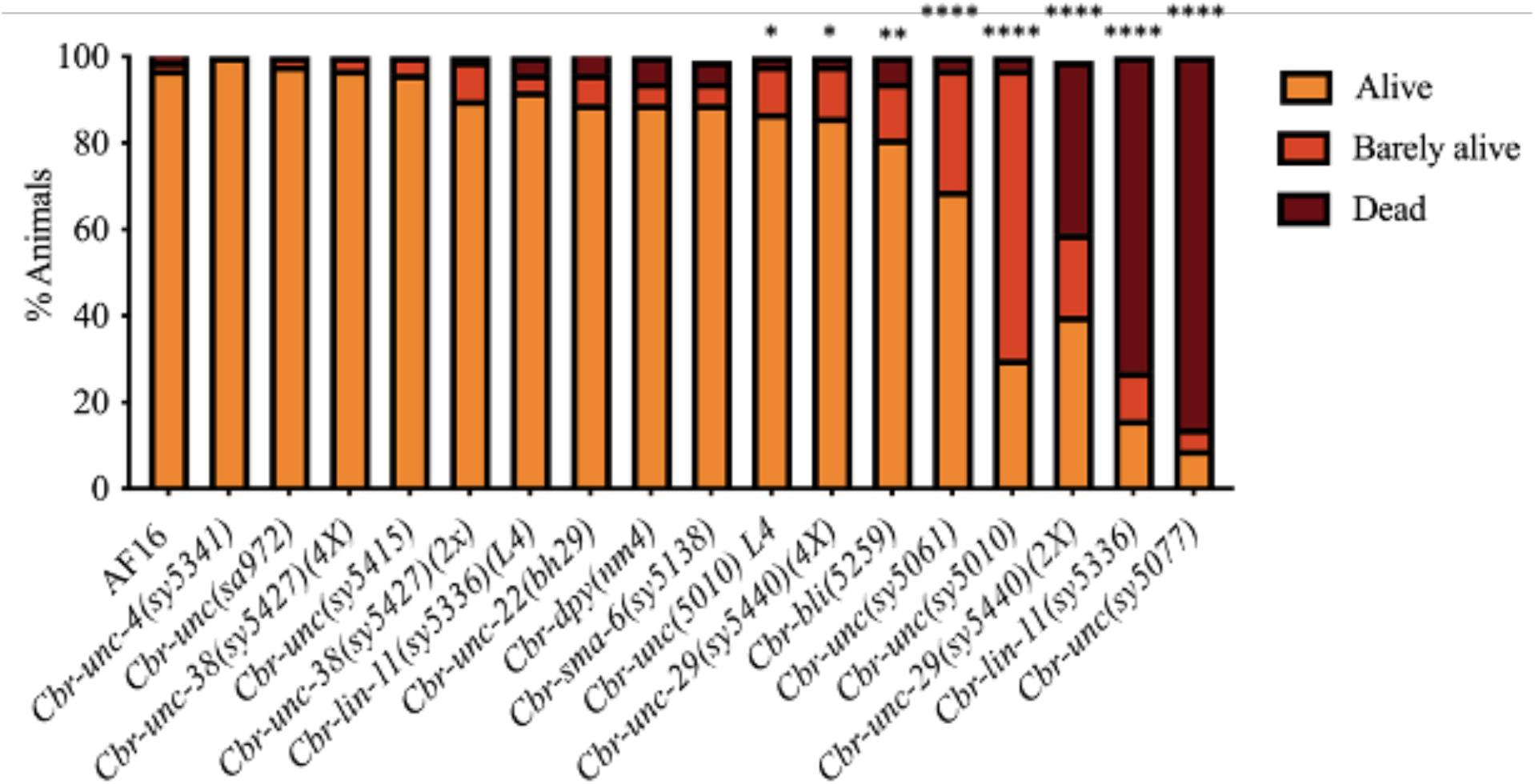
Heat shock responses of *C. briggsae* mutants. *C. briggsae* mutants belonging to *unc*, *sma*, *bli* and *lin* categories were given heat shock at 37^0^C for 2 hours. Based on their response to touch, the animals were classified into 3 categories after a 24-hour recovery period (see methods for detailed explanation on each category). While the 2x outcrossed *Cbr-unc-29(sy5440)* strain showed sensitivity to heat stress, the 4x strain was comparable to controls. In two cases, *Cbr-unc(sy5010)* and *Cbr-lin-11(sy5336)*, L4 stage larvae were also tested. The *Cbr-unc(sy5077)* animals are severely Unc and appear unhealthy, which likely affected their response to heat treatment. n = 47 - 152 worms collected from two to four batches for each of the strain. Statistical analysis was carried out using the chi-square test. *: (*p* < 0.05), **: (*p* < 0.01), ****: (*p* < 0.0001). Only those strains showing values significantly different from the AF16 control are indicated by stars.

## Supplementary tables

**Supplementary table 1.**
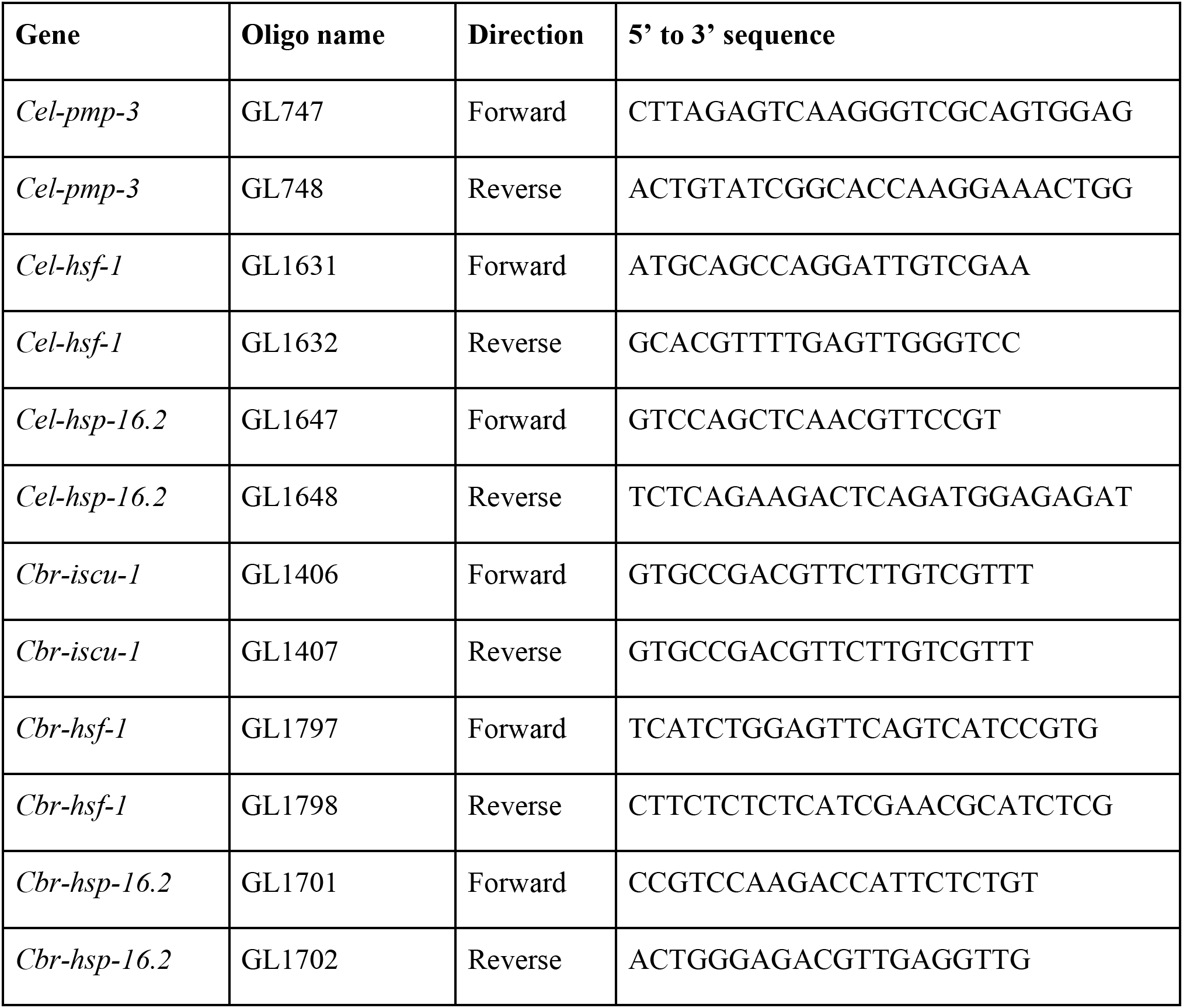
Primers used in qPCR experiments.

**Supplementary table 2.** Mean length and width of Caenorhabditis isolates. All values are in micrometers (µm). n, total number of animals combined from two batches. SD, Standard deviation.

| Strain | Length |  | Width |  | n |
| --- | --- | --- | --- | --- | --- |
|  | Average | SD | Average | SD |  |
| N2 hermaphrodite | 1051.34 | 81.40 | 61.54 | 2.58 | 10 |
| N2 male | 968.20 | 20.27 | 49.91 | 3.03 | 10 |
| AF16 hermaphrodite | 961.40 | 21.38 | 55.24 | 1.43 | 10 |
| AF16 male | 795.70 | 40.64 | 38.50 | 2.61 | 10 |
| HK104 hermaphrodite | 934.6 | 47.37 | 52.65 | 3.58 | 10 |
| HK104 male | 842.2 | 43.91 | 41.16 | 1.908 | 10 |
| VX34 hermaphrodite | 1076 | 93.15 | 54.55 | 2.079 | 13 |
| QX1410 hermaphrodite | 1023 | 94.23 | 55.67 | 3.55 | 16 |

## Notes

### Competing Interest Statement

The authors have declared no competing interest.

### Summary of Updates

Figure 1 has been revised by adding two new panels (C and D). A new supplementary table (#2) has also been included. These additions were followed by suitable modifications in Abstract, Methods, Results, and Discussion sections. Figure 1 legend has also been updated.

